# Metric Multidimensional Scaling for Large Single-Cell Data Sets using Neural Networks

**DOI:** 10.1101/2021.06.24.449725

**Authors:** Stefan Canzar, Van Hoan Do, Slobodan Jelić, Sören Laue, Domagoj Matijević, Tomislav Prusina

## Abstract

Metric multidimensional scaling is one of the classical methods for embedding data into low-dimensional Euclidean space. It creates the low-dimensional embedding by approximately preserving the pairwise distances between the input points. However, current state-of-the-art approaches only scale to a few thousand data points. For larger data sets such as those occurring in single-cell RNA sequencing experiments, the running time becomes prohibitively large and thus alternative methods such as PCA are widely used instead. Here, we propose a neural network based approach for solving the metric multidimensional scaling problem that is orders of magnitude faster than previous state-of-the-art approaches, and hence scales to data sets with up to a few million cells. At the same time, it provides a non-linear mapping between high- and low-dimensional space that can place previously unseen cells in the same embedding.

## 1 Introduction

Single-cell RNA sequencing (scRNA-seq) experiments provide quantitative measurements for thousands of genes across tens to hundreds of thousands or even millions of cells. The high-dimensionality as well as the sheer size of scRNA-seq data sets pose particular challenges for downstream analysis methods such as clustering and trajectory inference methods. An essential step in single-cell data processing is the reduction of data dimensionality to remove noise in gene expression measurements [27]. One of the most popular methods for dimensionality reduction of single-cell data is principle component analysis (PCA). PCA aims to maximize the variance in the reduced space and can be computed efficiently by a singular value decomposition. The existence of efficient implementations [1] has contributed to its routine application to large single-cell data sets.

Metric Multidimensional Scaling (MDS), on the other hand, aims to find an embedding that preserves pairwise distances between points (i.e. cells) which can improve the accuracy of various types of downstream analyses of single-cell data compared to PCA [27]. Its high computational cost, however, has hindered its wide application to single-cell data sets. In PHATE [20], for example, metric MDS is paired with a sampling-based approach to cope with its computational complexity.

Surprisingly, no algorithm is known that can solve metric MDS efficiently with more than a few thousand cells. Here, we provide the first such algorithm. Our contributions are two-fold: First, we provide a two-layer neural network approach that can solve the metric MDS problem for (single-cell) data sets with up to a few million data points (cells). This is orders of magnitude larger than current state-of-the-art methods can handle. Second, our approach for the first time learns a non-linear mapping of the high-dimensional points into the low-dimensional space, which can be used to place previously unseen cells in the same low-dimensional embedding.

### 1.1 Preliminaries

MDS comes in three different versions: 1) classical MDS, 2) metric MDS, and 3) non-metric MDS, aka ordinal scaling. While we focus here on metric MDS, it is important to understand all three methods and their differences.

Suppose we are given *n* data points *x_i_* ∈ ℝ^*m*^ that we want to embed into ℝ^*k*^ where *k < m*. Let *y_i_* be the corresponding point of *x_i_* in the low-dimensional space ℝ^*k*^.

#### Classical MDS

Classical MDS tries to map these data points into ℝ^*k*^ while trying to preserve the pairwise inner products ⟨*x_i_, x_j_*⟩. Specifically, it solves the optimization problem

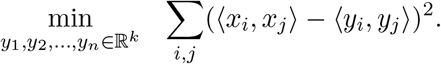

#### Metric MDS

Metric MDS tries to preserve the pairwise distances between the points, i.e., it solves the optimization problem

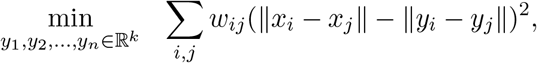

where ||.|| denotes the Euclidean norm and *w_ij_* ≥ 0 are some given weights.

#### Non-metric MDS

Non-metric MDS embeds the data into low-dimensional Euclidean space by preserving only the relative distance ordering, i.e, it solves the optimization problem

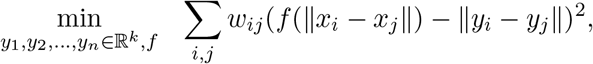

where *f* is a monotonically increasing function. Note, only the ordering of the pairwise distances is important here which should be preserved and not the actual distances.

All three formulations differ only in the objective functions that are minimized. While this seems like a minor difference, it has a substantial impact on its computability. It can be shown that classical MDS is equivalent to PCA when the input points are given explicitly and hence can be solved efficiently by a singular value decomposition. Thus, it can be solved efficiently for large data sets, having millions of cells. The computational complexity of metric MDS is fundamentally different. It has been shown that metric MDS is NP-hard when the target dimension is one [21] and it is believed that it is NP-hard in general. Hence, no efficient algorithm is likely to exist for solving this problem optimally. However, even finding a local minima is very time consuming. The most popular algorithm for solving this problem is the SMACOF algorithm by [8]. However, its running time grows quadratically in the number of data points *n*. It can only be applied to solve this problem with up to a few thousand data points.

There is one more important difference between classical MDS and metric MDS; it can be shown that in classical MDS the optimal solution corresponds to a linear mapping of the high-dimensional space ℝ^*n*^ into the low-dimensional Euclidean space ℝ^*k*^. This is not true for metric MDS. The optimal solution for metric MDS does not correspond to a linear mapping. Asking for a linear mapping leads to suboptimal solutions. Having a mapping from the input space ℝ^*m*^ into the output space ℝ^*k*^ is important for new, unseen data points. For instance, one can compute the mapping on a training set and apply the same mapping to the test set as it is common practice for other low-dimensional embedding and preprocessing methods like PCA. The SMACOF algorithm does not provide such a mapping.

Often, the input points are not given explicitly, but instead, their pairwise distances or pairwise scalar products are given. In this case, such a mapping cannot be provided.

### 1.2 Related work

Multidimensional scaling has a long history, see e.g., [13]. Classical MDS was first studied by [29] and independently by [12]. They used an eigenvector decomposition to solve the problem. Later, [18] defined the problem of metric MDS as an optimization problem and used a steepest descent approach for solving it. [8] improved the running time of Kruskal’s algorithm by using an iterative majorization approach. This algorithm is referred to as SMACOF algorithm. Surprisingly, it still represents the state-of-the-art for solving the metric MDS problem. The non-metric MDS problem was introduced by [18] and [14]. It is mainly used in the psychometric area.

The research on MDS can be split up into two branches; improving statistical performance and improving speed. Here, we focus on the latter one. The books by [5], and [2] provide an in-depth coverage on the statistical properties and applications of MDS. See also the book by [3] for a comparison of MDS to other embedding techniques.

When the input points are given explicitly, then classical MDS can be shown to be equivalent to PCA. Hence, computational speed is no issue in this case. When the input is instead a distance matrix, [22] and [33] used a divide-and-conquer approach for scaling up classical MDS to larger data sets. However, their approach only works for the classical MDS problem.

A technique called landmark MDS was introduced by [9]. The idea behind this approach is to select only a subset of the input points, called landmarks, compute the distance matrix between these points, and use only these points for the embedding. Hence, it can scale to larger data sets at the expense of ignoring the majority of the input points in the embedding process. This approach is also used in Isomap by [28]. [32] provided a similarity between kernel PCA and metric MDS. However, they are both similar but not equivalent due to their computational complexity (P vs. NP).

As already stated, classical MDS can be solved to optimality via a linear mapping. This is not true for the metric MDS. [24] set the weights *w_ij_* = 1/ ||*x_i_* − *x_j_*|| in the metric MDS problem and used a steepest descent algorithm for embedding the data. However, unlike the title suggests, it does not provide a mapping from the input space ℝ^*m*^ to the target space ℝ^*k*^.

There have also been some early attempts on solving the metric MDS problem using neural networks. This includes works by [19], and [23]. However, their approaches scaled to very small data sets with up to a few hundred data points only. While the neural network approach later was used for other non-linear embedding and dimensionality reduction methods and gave rise to autoencoders, see e.g., the seminal work by [15] they have never been used successfully for solving reasonably-sized metric MDS problems. [31] used a neural network approach for classical MDS and considered non-metric MDS in [30]. However, they did not consider the metric MDS problem.

## 2 Multidimensional scaling and linear mappings

In this section we will introduce projected metric MDS as an intermediate version of MDS that combines the optimization objective of metric MDS with the linear mapping obtained from classical MDS.

When applying MDS to a given input set, it is often important to also obtain the corresponding mapping, i.e., the mapping that maps the whole input space ℝ^*m*^ to the output space ℝ^*k*^. This is for instance necessary when MDS is used as a preprocessing step. One usually computes the embedding on the given training data set and applies the same mapping to the test data set, i.e., to new, unseen data points without recomputing the whole problem again. This is common practice in preprocessing steps, like PCA and also necessary in order to not induce a shift in the distribution of the test data. Hence, it is important to obtain such a mapping along with the actual embedding.

Let *X* ∈ ℝ^*n*×*m*^ be the matrix where the *i*th row is the *i*th input point *x_i_* ∈ ℝ^*m*^. We define the distance matrix *D_X_* ∈ ℝ^*n*×*n*^ as (*D_X_*)_*ij*_ = ||*x_i_* − *x_j_*||. Let 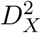 be the elementwise squared distance matrix, i.e., 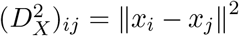. Let *Y* and *D_Y_* be defined accordingly for the output points *y_i_*.

Classical MDS minimizes the error on the pairwise scalar products, i.e., ∑_*i,j*_ (⟨*x_i_, x_j_*)−(*y_i_, y_j_*⟩)^2^. It has been shown that the optimal solution leads to a linear mapping which is obtained by the top-k eigenvectors of the Gram matrix *X*^⊤^*X*. The following computation shows that classical MDS can be translated into an optimization problem where approximately the error on the squared distances is minimized.

The objective function of classical MDS can be rewritten in matrix notation as

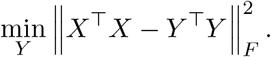

It holds that 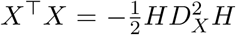, where 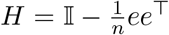 is a centering matrix with 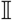 being the identity matrix and *e* ∈ ℝ^*n*^ the all-ones vector.

Hence, the objective function of classical MDS can be rewritten as

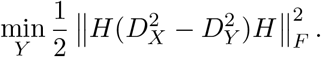

Note, this is very similar, however not equivalent to

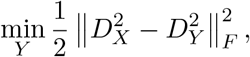

i.e., to the problem of minimizing the error on the squared distances. Compare this to the metric MDS problem that solves

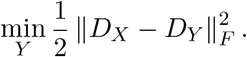

Hence, classical MDS can be seen as approximately minimizing the error of the squared distances while metric MDS tries to minimize the error of the distances. Thus, classical MDS can be a bit more sensitive to outliers in the data. One could then ask for an intermediate approach between classical and metric MDS; one could try to minimize the error on the distances while asking for a linear mapping. This gives rise to the projected metric MDS problem.

### Definition 1 (projected metric MDS).

*Given some input data X* ∈ ℝ^*n*×*m*^, *projected metric MDS solves the following optimization problem*

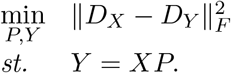

Matrix *P* ∈ ℝ^*m*×*d*^ along with the constraint forces the mapping to be linear. The following theorem relates the optimal solution of projected metric MDS to the optimal solution of metric MDS.

### Theorem 1.

*Let X* ℝ^*n*×*m*^ *be an input data matrix with centered rows, Y*\* ∈ ℝ^*n*×*k*^ *the solution of metric MDS and P*\* ∈ ℝ^*m*×*k*^ *the solution of the projected metric MDS. Then, the following inequality holds:*

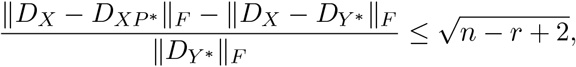

*where r* = rank(*X*).

In order to prove the theorem, we will need two technical lemmas (Lemma 1 and Lemma 2) and Proposition 1, where we show how to compute a projection matrix *P* such that the projected points are close to the given ones in a least-square sense. We show that this optimization problem has a closed formula solution and use this result as a way to generate a feasible solution for the projected metric MDS instance. After this we are ready to state the proof of Theorem 1..

### Definition 2.

*We say that rows of data matrix X are centered if*

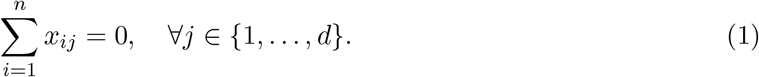

### Lemma 1.

*If X* ∈ ℝ^*n*×*m*^ *is centered, then for each P* ∈ ℝ^*m*×*k*^ *matrix XP is centered*.

*Proof.* Since *X* is centered, we have

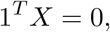

where 1 ∈ ℝ^*n*^ is vector of ones. From associativity of matrix multiplication we have that

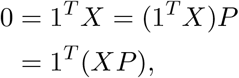

which implies that *XP* is centered.

### Lemma 2.

*Let X* ∈ ℝ^*n*×*m*^ *be a matrix with centered rows. Then*

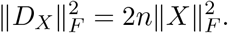

*Proof.* Since

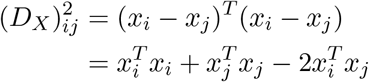

and rows of *X* are centered, we have

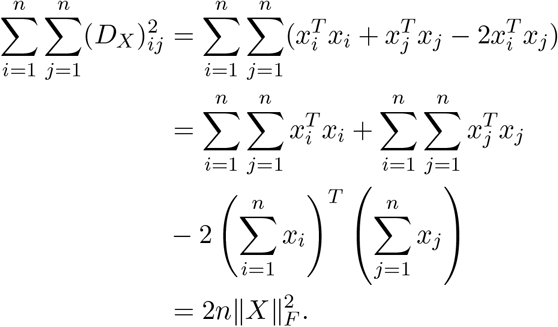

### Proposition 1.

*Let X* ℝ^*n*×*m*^ *be an input data matrix,* 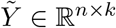 *for some k < m, and a singular value decomposition of X given by the following:*

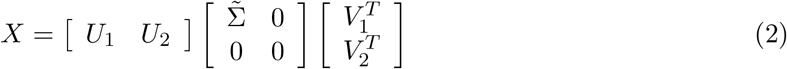

*where U* = [*U*_1_ *U*_2_]*, U*_1_ ∈ ℝ^*n*×*r*^, *U*_2_ ∈ ℝ^*n*×(*n*−*r*)^ *and V* = [*V*_1_ *V*_2_]*, V*_1_ ∈ ℝ^*m*×*r*^, *V*_2_ ∈ ℝ^*m*×(*m*−*r*)^ *are orthogonal matrices.*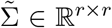 *is a diagonal matrix that contains r non-zero singular values such that σ*_1_ ≥ *σ*_2_ ≥ *…. σ_r_ >* 0, *where r is a rank of matrix X*.

*The solution of the following optimization problem*

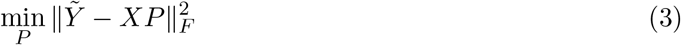

*is given by*

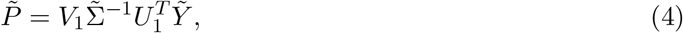

*whose objective value is*

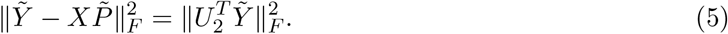

*Proof*. The objective function in (3) can be rewritten as follows:

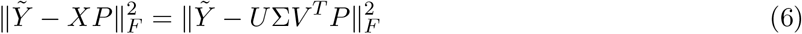

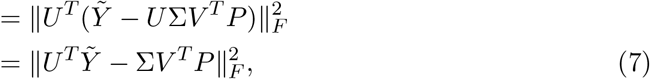

where last equality follows from unitary invariance of Frobenius norm. We introduce substitution

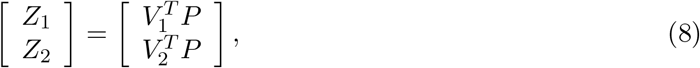

such that the objective in (7) is

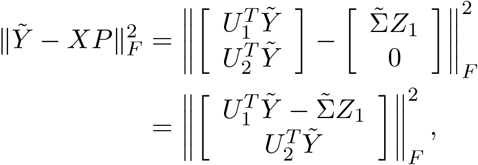

where 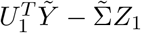 can be set to zero matrix when *Z*_1_ is a solution of the following equation:

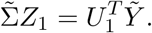

Indeed, 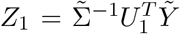, while *Z*_2_ can be set to 0 since it does not affect the objective value.

Substitution of *Z*_1_ from (8) gives the solution (4) whose objective value is given in (5).

Now we are ready to state the proof of Theorem 1

*Proof of Theorem 1.* In order to prove the inequality, we first bound 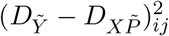.

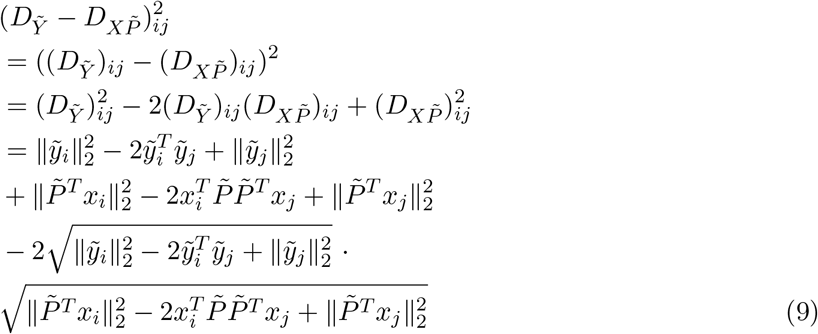

Now, from Cauchy-Schwartz inequality follows that 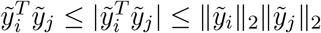 which gives

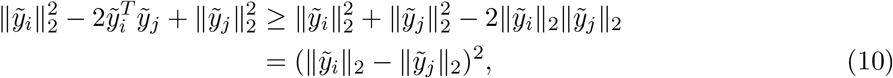

and analogously

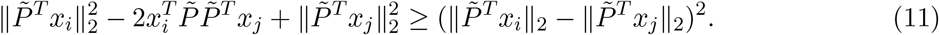

From (9), (10), and (11) follows that

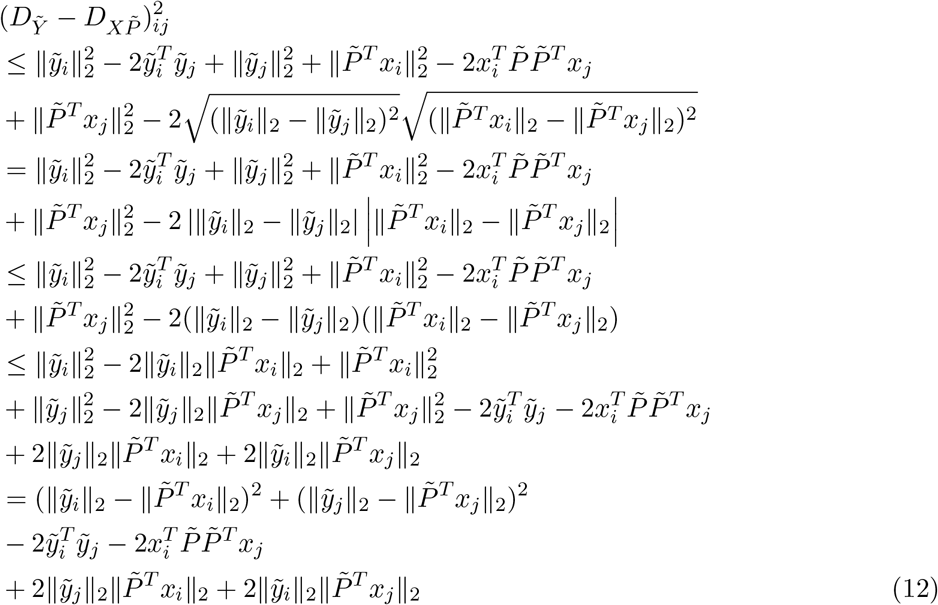

which after summing up both sides gives

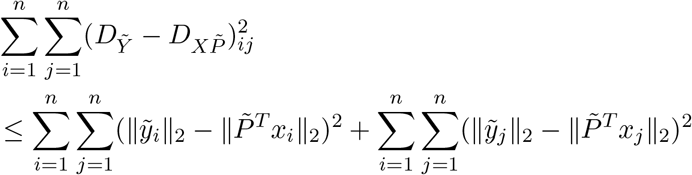

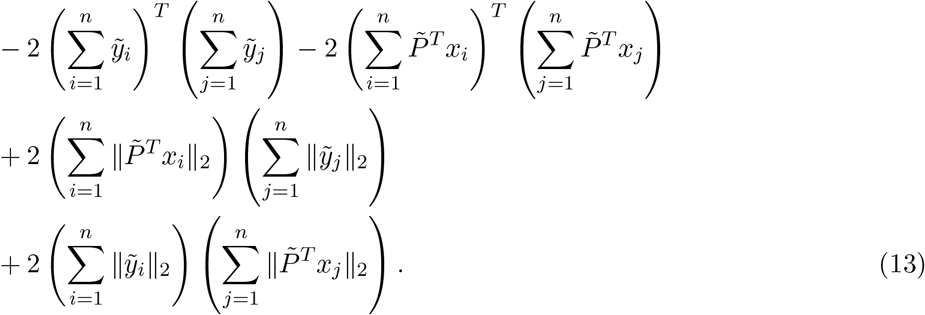

Now, from | ||*x*|| − ||*y*|| | ≤ ||*x* − *y*|| and Lemma 1 we obtain the following bound

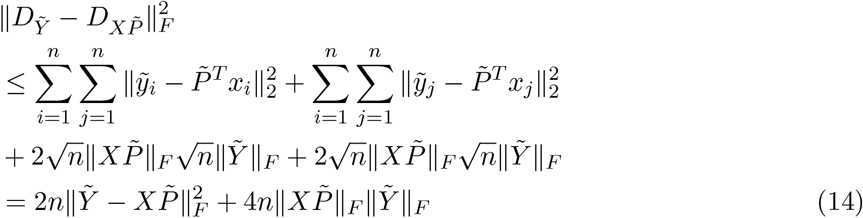

Now, from Proposition 1 we conclude that

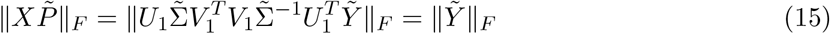

and

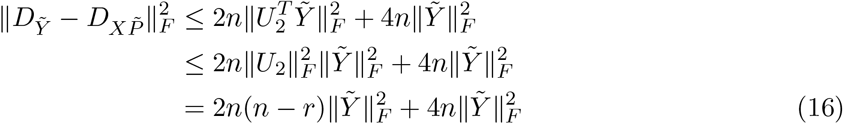

Inequality (16) and Lemma 2 imply

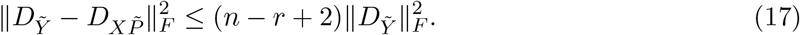

Finally, from triangle inequality and (16) we conclude that

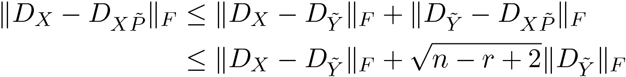

which completes the proof.

While the theorem proves that the objective function values of projected metric MDS and metric MDS do not differ too much, in practice they still might provide substantially different solutions. See Figure 1 for an example. Hence, we conclude that it is really necessary to have a nonlinear mapping between the input and the output space. This motivates the neural network approach introduced below.

**Figure 1:**
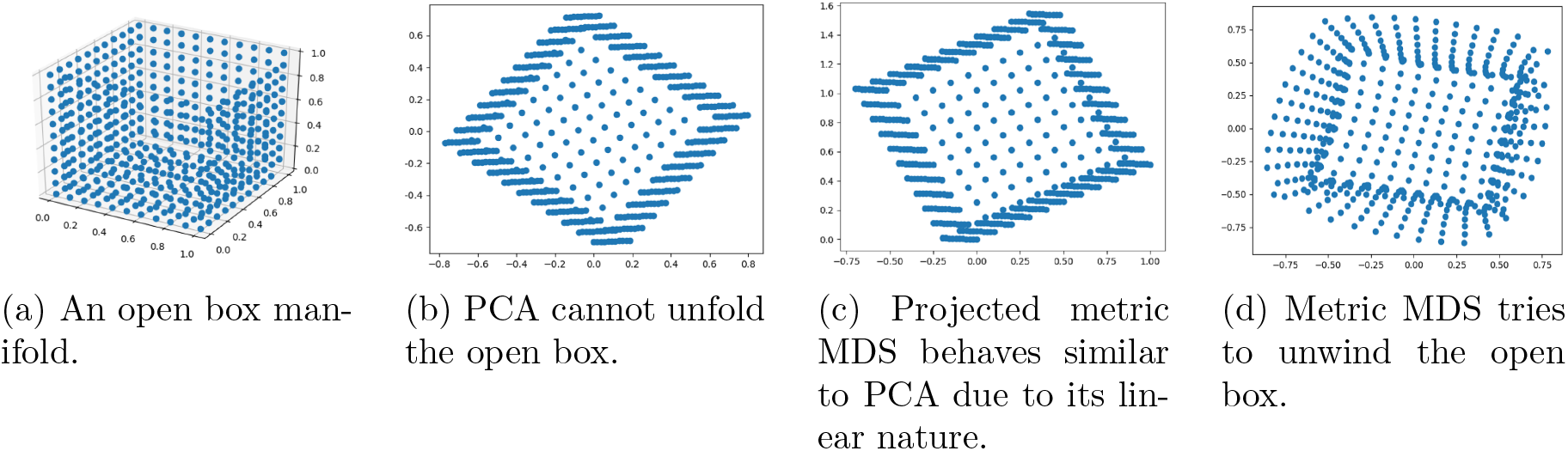
We use an open box example (a) in order to illustrate the power of nonlinear mapping, such as metric MDS (d), over the linear mapping, such as PCA (b) and projected metric MDS (c).

## 3 The neural network approach

Here we will describe the neural network architecture that we used as well as the algorithm for computing the metric MDS mapping. We use a very simple fully-connected neural network with a single hidden layer and a tanh activation function. The size of the input layer corresponds to the dimension of the input data *n*, while the size of the output layer corresponds to the dimension *k* of the output data. The size of the hidden layer is chosen as an estimate of the intrinsic dimension of the input data set. In practice, we estimated the intrinsic dimension by computing a SVD of the input data and selecting the top *k* singular values that retain at least 95% of the data variance.

The motivation for our simple neural network architecture comes from the fact that any feed-forward neural network with only a single hidden layer and any infinitely differentiable sigmoidal activation function, i.e., a function that retains the S shape can uniformly approximate any continuous function on a compact set, see, e.g., [6]. Furthermore, any mapping from the input points to the output points can be extended to a continuous mapping from ℝ^*n*^ to ℝ^*k*^, provided that the set of input points is finite and no two input points share the same coordinates, i.e., *x_i_* ≠ *x_j_* for *i* ≠ *j*. Hence, a neural network should be able to approximate that mapping arbitrarily well. Due to the simplicity, differentiability, and the small number of parameters compared to the number of input points and features, there is little chance of overfitting. As can be seen in the experiments, the found mapping generalizes well on test data.

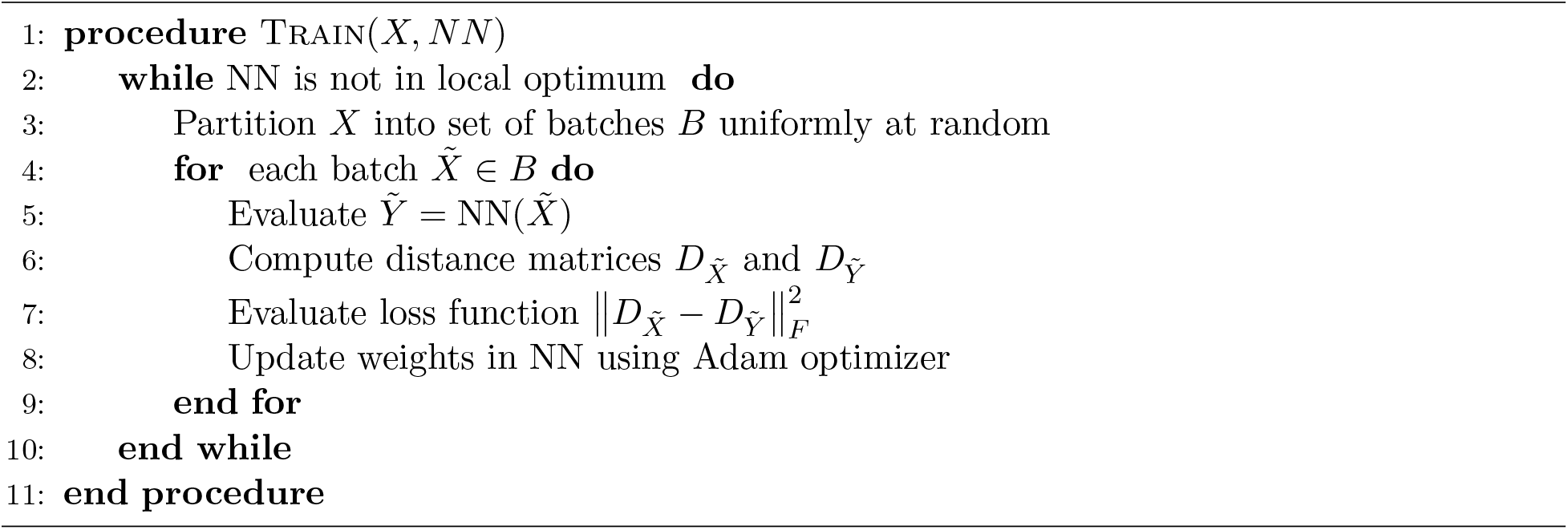
Algorithm 1. NN approach for metric MDS problem

We adopt the standard batch stochastic gradient descent approach and partition the input data set into a set of batches. We optimize the weights in our simple neural network following the idea of the Siamese neural network approach, see, e.g., [4]. The basic idea of this approach is that in each learning step the neural net is shown two points, say *x_i_* and *x_j_*. The outputs are stored for both points, say *y_i_* and *y_j_* respectively and the distance ||*y_i_* − *y_j_*|| between the output vectors are calculated. A loss function is defined in terms of the squared difference of this distance and the distance between the points in the input space ||*x_i_* − *x_j_*||, for all pairs of input points in the current batch. The weights of the neural network are updated using the Adam optimizer. Since the data is shuffled after each epoch, we sample the input distance matrix uniformly at random which provides an unbiased estimate of it. We set the batch size to 256 points which provides a good tradeoff between memory consumption and number of iterations needed to converge. All the steps of our simple but efficient approach are summarized in Algorithm 1.

## 4 Experiments

We implemented the neural network (NN) approach and compared it to other methods for solving MDS^1^. Specifically, we compared it to the SMACOF algorithm which still represents the state-of-the-art for solving the metric MDS problem. However, due to its inherent quadratic runtime and space complexity, it is prohibitive to run it on data sets with more than a few thousand data points. To still assess the quality of our neural network approach, we also compared it to a random projection (RP) approach. In the random projection approach, a random Gaussian matrix is used for projecting the points into a low-dimensional Euclidean space. The well-known Johnson-Lindenstrauss lemma [17] states that such a random projection can embed any *n*-point data set into a low-dimensional Euclidean space of dimension at most 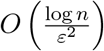 while incurring a multiplicative distortion error of no more than 1 + *ε* for any small *ε* > 0 in the worst case. [7] showed that in general this approach works surprisingly well when trying to embed a data set while preserving inter-cluster distances. We also compared our neural network approach to PCA to demonstrate that the preservation of distances as explicitly optimized for in mMDS cannot be obtained as a by-product of a much simpler optimization model (here classical MDS). Finally, we also compared it to the projected metric MDS problem that we have defined in Section 2. We solved the projected metric MDS problem using a quasi-Newton method combined with a smoothing technique. Note that this approach also does not scale well to large data sets.

We ran the experiments on a Ryzen 9 3900X CPU with 12 cores running at 3.8 GHz, 64 GB DDR4 RAM and a RTX 2080 graphics card using an Ubuntu 19.10. operating system. All implementations were done in Python 3.7. We used PyTorch 1.5.1 for the neural network approach. We used the implementation of the random projection, PCA, and SMACOF algorithm from scikit-learn 0.22.1.

### 4.1 Comparison of loss and running time of metric MDS

Figure 2 shows the metric MDS loss achieved by the different methods on the USPS, MNIST, CIFAR10, and SVHN data sets for different embedding dimension *k*. For each data set, we computed the embedding on the train data set and then applied the mapping of the input space to the lower-dimensional output space provided by each method (except SMACOF) to new, unseed data points from the test data set. Since the SMACOF algorithm does not provide such a mapping we recomputed the embedding of combined train data and test data and reported the loss of SMACOF on the test data set in Figure 2b. On the smallest USPS data set our neural network approach is typically at least as good as the SMACOF algorithm, which we could not run on other (larger) data sets due to its prohibitive running time. As expected, both methods yielded substantially smaller loss across data sets than the remaining methods that do not explicitly optimize the same objective. Note that the metric MDS loss function is plotted on a logarithmic scale.

**Figure 2:**
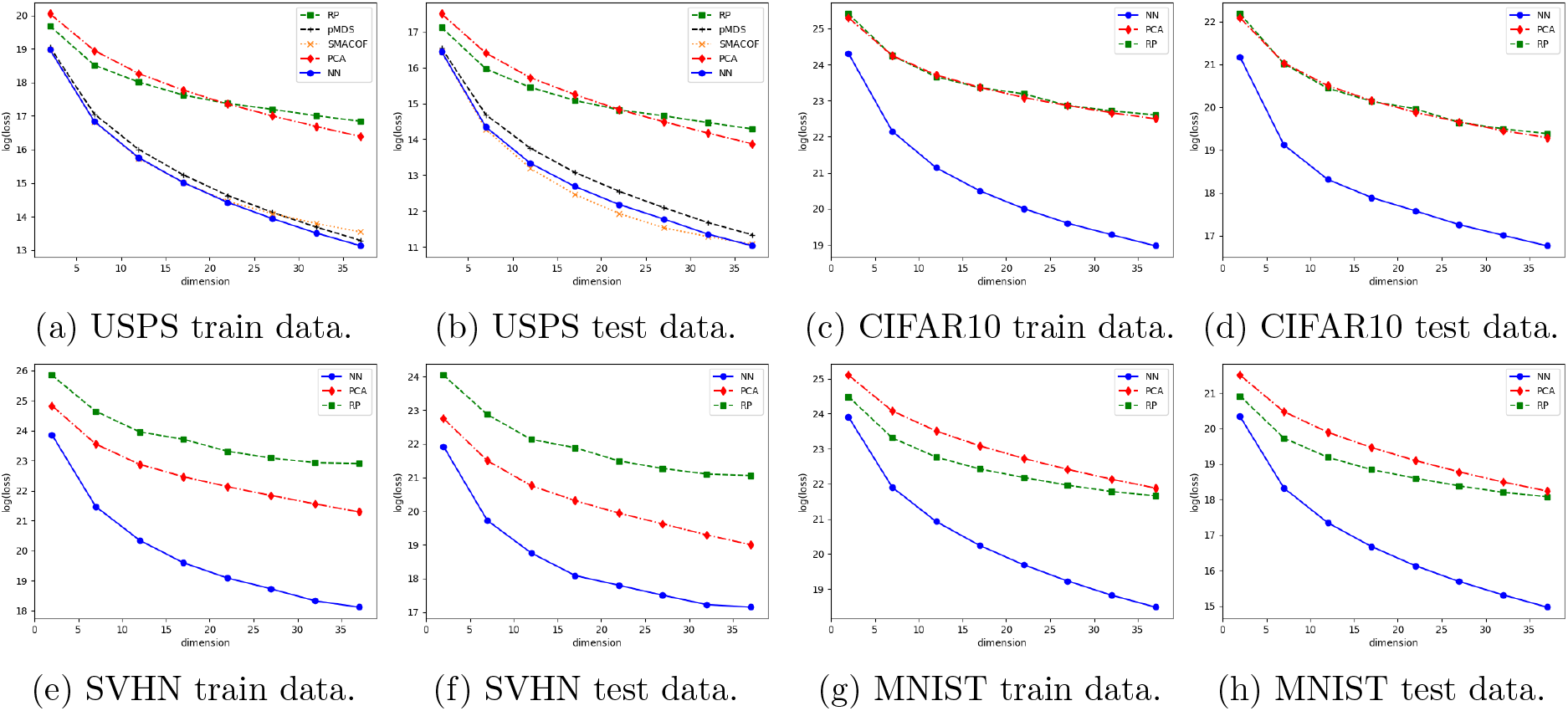
The loss of the metric MDS problem for different values of the target dimension for train and test data sets. The loss function is displayed on a logarithmic scale. Due to its quadratic running time, SMACOF was run only on the smallest USPS data set.

Figure 3 shows consistent results on three word embedding data sets that represent words as vectors that geometrically capture the semantics of the words. More precisely, the Glove data set consists of 2 data sets: `glove.6B` learned word vectors on Wikipedia 2014 dump (size 400K), and `glove.840B` learned word vectors on Common Crawl corpus (size 2.2.M). The FastText data set `wiki-news-300d-1M` holds 1M word vectors trained on Wikipedia 2017 dump, UMBC webbase corpus and statml.org news dataset. We had to exclude SMACOF from this comparison due to its quadratic running time.

**Figure 3:**
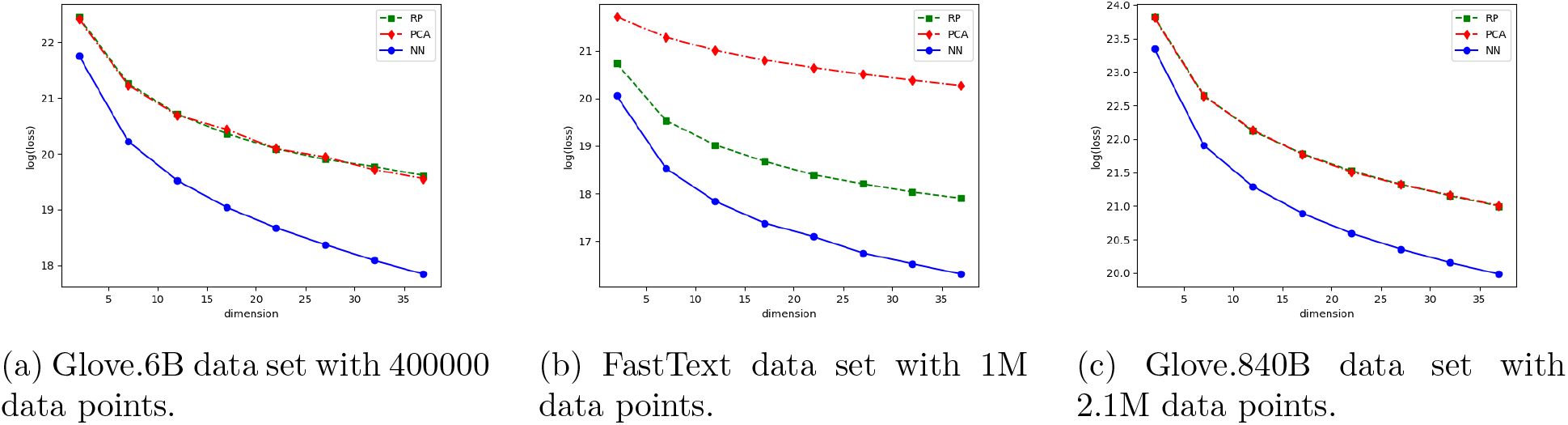
Comparison of the loss function of the metric MDS problem for random projections (RP), PCA, and our neural network (NN) approach.

We also report the running time of our approach in Table 1 and compare it to the running time of the SMACOF algorithm when embedding the different data sets into dimension *k* = 12. We observed that neither the running time of SMACOF nor of our approach depends on the dimension *k*. Since the SMACOF algorithm can handle only small data sets we subsampled the MNIST data set. We always ran our approach for 1000 epochs on all data sets. Table 1 shows that the running time of the SMACOF algorithm grows quadratically in the number of data points. In contrast, our approach shows an approximately linear dependence, which allows it to be applied to large data sets where it is orders of magnitude faster than the current state-of-the-art approach.

**Table 1:** Running times of SMACOF and our neural network based approach on data sets of different size. The dimension of the output space *k* was fixed to 12 in this experiment. Times are presented in format hh:mm:ss. Missing values indicate that the solver did not finish within 24 hours.

| Dataset | size | n | SMACOF | NN |
| --- | --- | --- | --- | --- |
| USPS | 7291 | 256 | 0:11:28 | 0:05:14 |
| MNIST_5 | 5000 | 784 | 0:21:41 | 0:03:43 |
| MNIST_10 | 10000 | 784 | 1:22:12 | 0:04:09 |
| MNIST_15 | 15000 | 784 | 3:05:47 | 0:05:47 |
| MNIST_20 | 20000 | 784 | 5:38:38 | 0:07:11 |
| MNIST_25 | 25000 | 784 | 8:47:28 | 0:08:53 |
| MNIST_30 | 30000 | 784 | 12:40:55 | 0:10:20 |
| MNIST | 60000 | 784 | - | 0:21:21 |
| CIFAR10 | 50000 | 3072 | - | 0:27:45 |
| SVHN | 73257 | 3072 | - | 0:50:56 |
| FastText | 999994 | 300 | - | 2:52:18 |
| Glove.6B | 400000 | 100 | - | 1:02:32 |
| Glove.840B | 2196016 | 300 | - | 6:07:24 |

## 5 Metric MDS based clustering of scRNA-seq data

In this section we demonstrate the utility of metric MDS for the clustering of single-cell RNA sequencing (scRNA-seq) data. The unsupervised clustering of scRNA-seq data allows to identify known as well as novel cell types based on the cell’s transcriptomes. Seurat [25] is the most widely used computational method for clustering of scRNA-seq. It is based on the Louvain clustering algorithm and relies on a prior preprocessing of the data that includes, among others, a dimensionality reduction step using principle component analysis (PCA). A major advantage of metric MDS over PCA is its flexibility with respect to the distance metric that is used in the underlying optimization problem. We therefore compared the standard Seurat clustering pipeline to a pipeline in which we replaced the PCA step by our metric MDS approach but kept all other computational steps identical. In metric MDS we experimented with 3 different distance metrics, the Euclidean, cosine, and correlation based distance.

We compared the PCA and metric MDS based clustering approaches on all but one real data sets that were used in [10] to benchmark clustering methods using cell phenotypes defined independently of scRNA-seq. Following [10], we labeled cell types based on FACS sorting in the Koh data set, and grouped cells according to genetic perturbation and growth medium in the Kumar data set. In data set Zhengmix4eq (Zhengmix4uneq), the authors in [10] randomly mixed equal (unequal) proportions of presorted B cells, naive cytotoxic T cells, CD14 monocytes, and regulatory T cells. Data set Zhengmix8eq additionally included equal proportions of CD56 NK cells, naive T cells, memory T cells, and CD4 T helper cells. We excluded a single data set in which ground truth labels correspond to collection time points that all methods in [10] failed to reconstruct. In agreement with recent clustering benchmarks [10, 11] we used the Adjusted Rand Index (ARI) [16] and Normalized Mutual Information (NMI) [26] to quantify the similarity of inferred to ground truth clusterings.

On average, metric MDS yielded more accurate clusterings than when applying PCA, independent of the specific distance metric used (Table 2). Clusterings obtained from embeddings computed by metric MDS using correlation or cosine based distances were most accurate, and achieved a substantial improvement compared to PCA on the three most difficult (with respect to PCA performance) data sets (Figure 4). Performance metric NMI provided a consistent picture of method performance. A t-SNE visualization of the two embeddings highlights the better separation of cell types by metric MDS compared to PCA on data set Zhengmix4eq (Figure 5), especially between naive cytotoxic and regulatory T cells.

**Table 2:** Average scores of ARI and NMI metrics across 5 real data sets. Clusterings are obtained from embeddings computed by mMDS using correlation, cosine and euclidean distance.

| Metrics | correlation | cosine | euclidean | Seurat (PCA) |
| --- | --- | --- | --- | --- |
| ARI | 0.9177 | 0.9179 | 0.8681 | 0.8180 |
| NMI | 0.9325 | 0.9302 | 0.8880 | 0.8757 |

**Figure 4:**
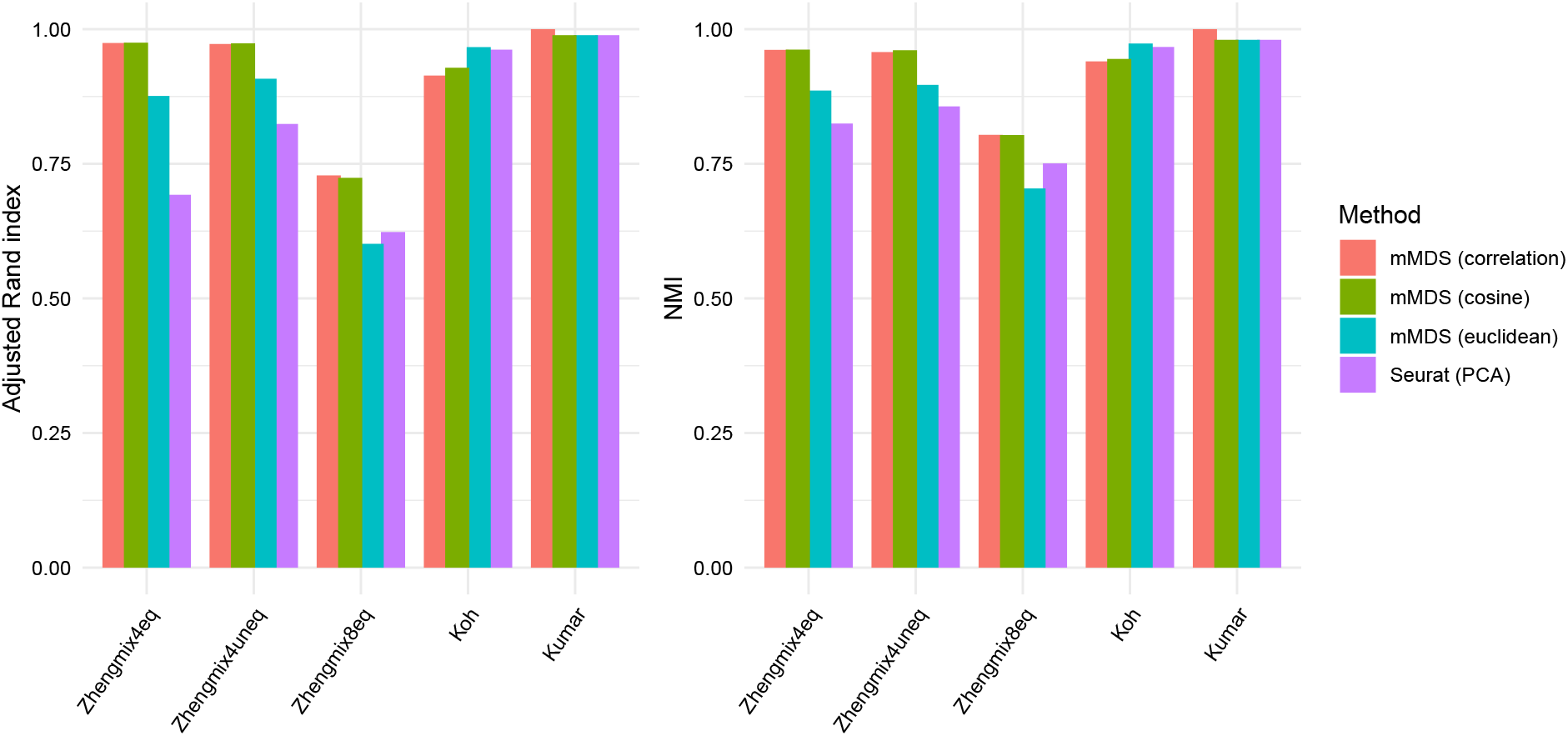
Comparison of metric MDS using different distance metrics and PCA in scRNA-seq clustering on 5 real data sets.

**Figure 5:**
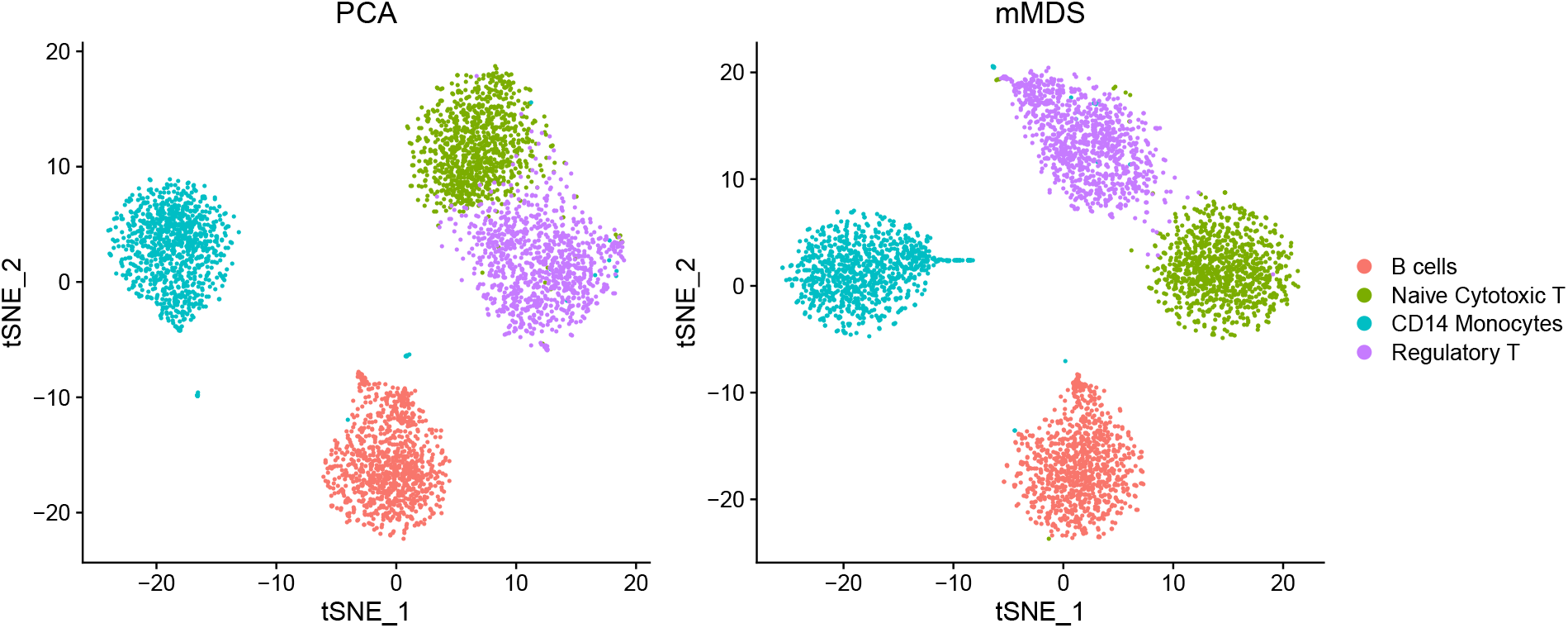
t-SNE visualizations of the embeddings computed by PCA (left) and correlation based metric MDS (right) on Zhengmix4eq data set.

## 6 Conclusion

We presented a two-layer neural network approach for solving the metric multidimensional scaling problem. Our approach provides two advantages over previous state-of-the-art approaches; it is orders of magnitude faster and scales to much larger data sets with up to a few million data points and may thus represent a viable alternative to the widely used PCA in single-cell analysis. At the same time it provides a mapping of the input space to the output space. This allows to apply the same embedding to new, unseen data, which prevents inducing a shift in the data distribution for test data.

## Footnotes

1 The source code is publicly available via https://github.com/dmatijev/nnMDS.git

